# EthoClaw: An Integrated AI Workflow Platform for Automated Analysis in Neuroethology

**DOI:** 10.64898/2026.03.25.714141

**Authors:** Ke Chen, Ziming Chen, Dagang Zheng, Xiang Fang, Jinghong Liang, Zhenyong Li, Yufeng Chen, Jiemeng Zou, Bingdong Cai, Shanda Chen, Kang Huang

## Abstract

Computational methods have advanced the analysis of animal behavior, yet significant challenges remain in data standardization, analytical reproducibility, and workflow integration. Existing computational solutions often demand extensive programming proficiency or compel users to navigate a highly fragmented ecosystem of disconnected tools for tracking, statistical analysis, and visualization. Here, we present EthoClaw, an open-source, artificial intelligence-driven workflow platform built upon the OpenClaw agentic framework, functioning as a locally deployable AI assistant for behavioral research. EthoClaw provides an integrated computational infrastructure that seamlessly bridges the gap between raw behavioral video acquisitions and publishable scientific results. In this study, we demonstrate the platform’s capacity to natively ingest video data via a dual-mode tracking architecture: utilizing ultra-fast image processing for rapid object detection, and leveraging the SuperAnimal methods for precise, markerless postural tracking. To ensure maximal interoperability, EthoClaw automatically converts various tracking data formats into DeepLabCut-compatible formats, enabling high-throughput phenotyping by generating publication-quality visualizations alongside rigorous multidimensional statistical profiling. Furthermore, the platform incorporates a large language model (LLM)-driven reporting module that dynamically synthesizes analytical documents, ensuring methodological transparency. Through an open field test, we validate the practical usability of EthoClaw while accelerating computational throughput by localizing heavy video processing to circumvent cloud bandwidth bottlenecks. Operating via an omnichannel natural language interface that integrates seamlessly with ubiquitous instant messaging software, EthoClaw democratizes advanced computational behavioral analysis, offering a holistic, highly efficient ecosystem that enforces experimental reproducibility and open science principles.

## Introduction

Animal behavior analysis currently stands at a critical technological juncture, where traditional observational paradigms are being rapidly augmented by unprecedented computational capabilities. Over the past decade, advances in computer vision and deep learning have enabled researchers to capture high-fidelity kinematic data from single experimental sessions (Pereira et al., 2020; Mathis et al., 2018). These high-dimensional datasets hold immense potential for deciphering complex behavioral motifs and identifying neural correlates of behavior. However, translating raw experimental video into biologically meaningful insights remains a highly fragmented and technically prohibitive process.

The prevailing analytical workflow typically involves a disjointed sequence of operations: local computational environment configuration, heterogeneous data format matching, video acquisition, pose estimation, feature extraction, statistical testing, and final visualization. Each stage inherently relies on distinct software architectures and varying levels of analytical expertise. Consequently, researchers frequently resort to developing custom scripts to bridge these methodological gaps. More critically, the absence of standardized data pipelines and unified analytical protocols severely undermines experimental reproducibility, exacerbating the broader reproducibility crisis in the life sciences (Baker, 2016).

While several computational tools have emerged to address specific bottlenecks, a unified solution remains elusive. Prominent frameworks such as DeepLabCut (Mathis et al., 2018) and SLEAP (Pereira et al., 2022) provide state-of-the-art pose estimation, which has been further enhanced by recent transformer-based architectures designed to mitigate tracking drift across diverse species (Tang et al., 2025). Beyond foundational 2D tracking, recent methodological advancements have successfully pushed the field toward complex 3D markerless motion capture (Huang et al., 2021; Han et al., 2022), sophisticated multi-animal social interaction mapping (Han et al., 2024), behavioral modeling of free social interactions (Han et al., 2026a), and explorations of the relationship between hierarchical behaviors and the brain (Han et al., 2026b). However, despite these remarkable modular achievements, these tools primarily isolate their focus to specialized tracking or specific analytical phases. Integrated pipelines like ezTrack (Pennington et al., 2019) offer robust analysis but are generally constrained to highly specific paradigms. Commercial suites provide end-to-end functionality but incur prohibitive costs and operate as proprietary black boxes, limiting methodological transparency.

To address this methodological gap, we developed EthoClaw, a domain-specific computational ecosystem tailored for behavioral neuroscience. Built upon the OpenClaw agent framework, EthoClaw operates as a locally deployable AI research assistant. By keeping computation-heavy tasks on local hardware while utilizing cloud-based LLM APIs merely for lightweight workflow orchestration, EthoClaw overcomes the severe bandwidth limitations and latency issues associated with uploading terabytes of raw experimental video to the cloud. The platform introduces a dual-mode tracking system—combining classical computer vision for rapid object detection with a deep learning model for intricate postural analysis. Furthermore, it ensures widespread interoperability by harmonizing diverse data formats into DeepLabCut-compatible schemas. In this paper, we present the capabilities of EthoClaw, evaluate its computational robustness, and demonstrate its utility in streamlining behavioral research workflows while enforcing rigorous open-science standards.

## Results

### An Agentic Workflow Ecosystem for Behavioral Phenotyping

The computational paradigm for behavioral phenotyping has evolved rapidly. Historically, researchers relied on rigid, task-specific software—operated via traditional graphical user interfaces (GUIs) or command-line interfaces—demanding extensive manual configuration. The recent advent of online Large Language Models (LLMs) introduced natural language processing to scientific workflows, yet these foundational models lacked the capacity to directly execute complex, multi-step analytical pipelines. This limitation catalyzed the development of online, cloud-based AI agents capable of autonomous tool-use. However, fully cloud-dependent agentic systems present significant bandwidth and latency issues for laboratories handling massive experimental video datasets, as uploading terabytes of raw video for cloud-based tracking is highly impractical.

To bridge this gap, we sought to establish a computational architecture that minimizes technical barriers while ensuring rapid processing of high-dimensional local data. We developed EthoClaw atop the OpenClaw agentic framework, engineered specifically as a locally deployable agent that executes on laboratory hardware (Fig. 1). By keeping heavy computational tasks—such as video ingestion, image processing, and deep learning inference—strictly local, EthoClaw circumvents the latency of cloud data transfer while utilizing API-driven LLMs purely for lightweight workflow orchestration and code generation. This architecture fundamentally transitions user interaction to conversational natural language dialogue. Inheriting OpenClaw’s robust omnichannel capabilities, EthoClaw connects directly to popular instant messaging software. This integration allows researchers to conveniently orchestrate complex analytical pipelines, initialize computationally intensive tracking tasks on their local GPUs, and receive real-time analytical reports directly within their daily communication channels. The architecture also establishes a seamless integration with third-party plugin marketplaces like Clawhub, ensuring the platform’s extensibility for future ethological modeling.

**Fig. 1.**
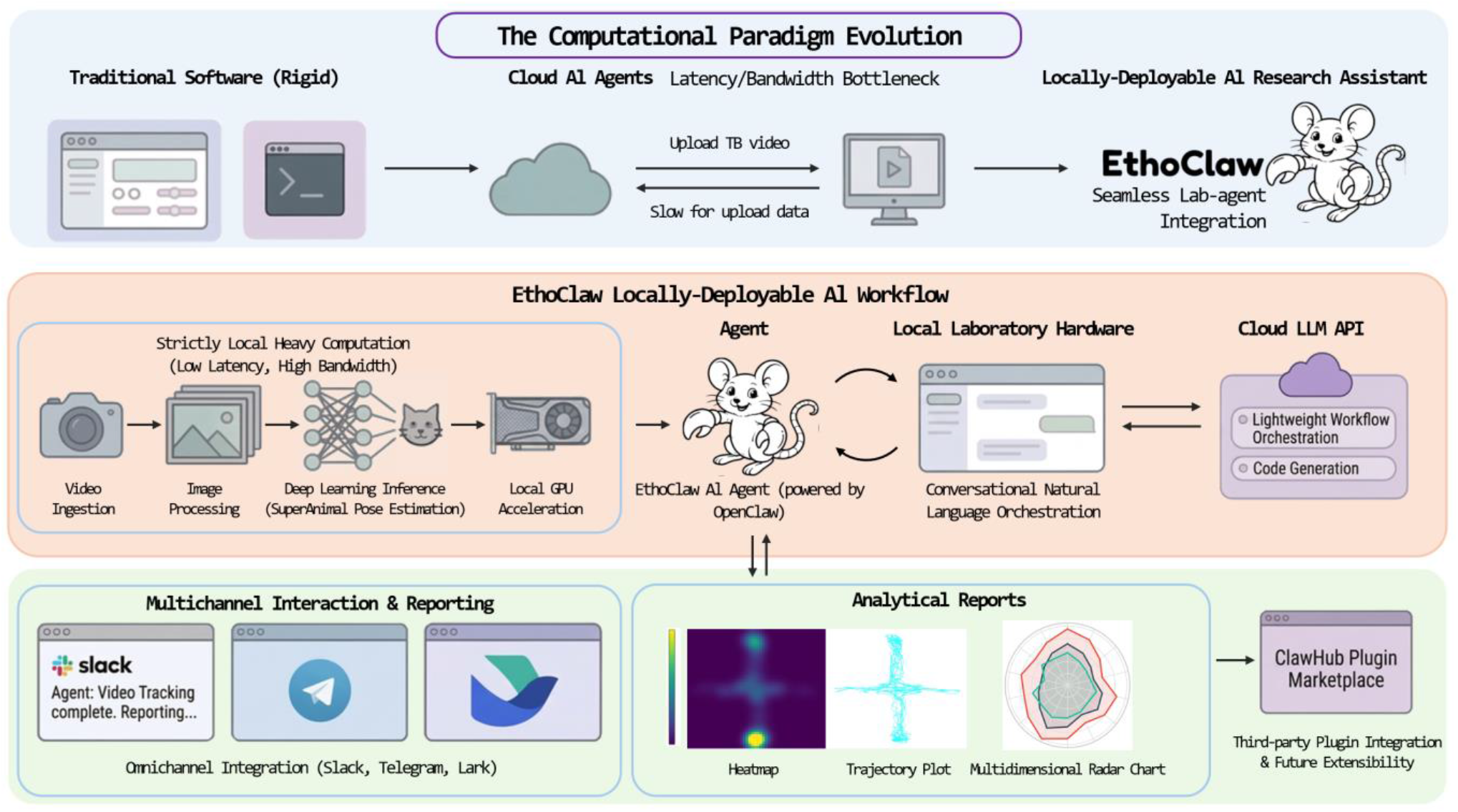
System architecture of EthoClaw and the evolution of computational paradigms in behavioral phenotyping. Evolution of computational paradigms toward locally-deployable AI assistants. EthoClaw’s hybrid workflow: computation-heavy video processing and pose estimation are kept strictly local to avoid bandwidth bottlenecks, while cloud LLMs are utilized solely for lightweight natural language orchestration. Omnichannel integration with standard instant messaging apps for accessible conversational interaction, real-time analytical reporting, and third-party plugin extensions via ClawHub.

### Dual-Mode Object Detection and Pose Estimation

A major bottleneck in behavioral analysis is the transition from raw video to coordinate data. We evaluated EthoClaw’s capacity to natively ingest and process raw experimental videos without relying on intermediate software. By employing adaptive computer vision algorithms to autonomously identify animal subjects, EthoClaw introduces a highly flexible dual-mode tracking architecture (Fig. 2 top).

**Fig. 2.**
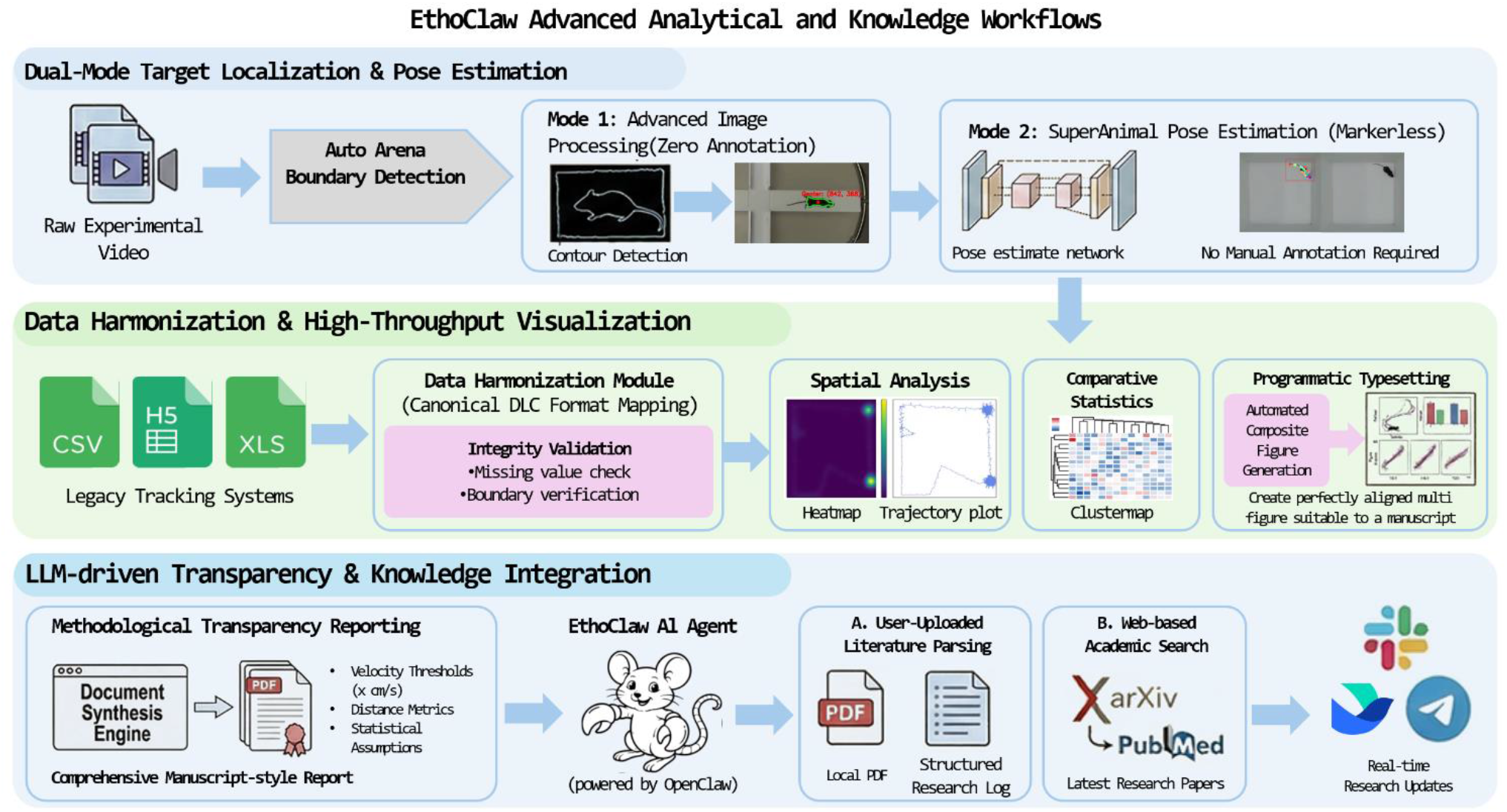
EthoClaw Advanced Analytical and Knowledge Workflows. Dual-Mode Object Detection and Pose Estimation: EthoClaw natively ingests raw video and detects arena boundaries. It features a dual-mode tracking architecture: Mode 1 uses zero-annotation image processing to rapidly extract centroid coordinates, while Mode 2 leverages the SuperAnimal deep learning methods for precise, markerless multi-node pose estimation, eliminating manual frame annotation. Data Harmonization and High-Throughput Visualization: The Data Harmonization Module parses heterogeneous tracking outputs into a canonical DeepLabCut (DLC)-compatible format, performing integrity validations like missing value checks. Using this standardized data, the engine automates high-throughput visualizations and programmatically aggregates them into publication-ready composite figures. LLM-driven Transparency and Knowledge Integration: To ensure methodological transparency, the Document Synthesis Engine automatically generates manuscript-style HTML/PDF reports detailing exact analytical parameters. Additionally, the Knowledge Integration Pipeline acts as a research assistant by (A) parsing user-uploaded PDFs into structured logs and (B) executing automated API searches to deliver real-time literature updates directly via instant messaging.

First, it features an advanced image processing-based object detection pipeline. While slightly less robust than deep neural networks in complex environments, this classical computer vision approach requires zero data annotation and boasts an extremely low computational footprint. Crucially, it executes with exceptional speed—often achieving faster-than-real-time processing rates—relying entirely on standard CPU hardware without the necessity for expensive GPU accelerators. This hardware accessibility and rapid processing capability render it an optimal, highly efficient choice for high-throughput locomotor paradigms characterized by clean backgrounds and high subject-to-background contrast.

For more granular postural analyses, EthoClaw interfaces directly with the SuperAnimal method from the DeepLabCut Model Zoo, specifically deploying the pre-trained superanimal_topviewmouse model. We successfully demonstrated the precise, markerless tracking of distinct anatomical nodes in black mice within standard 2D top-down assays, such as the Open Field Test (OFT). During inference, the model robustly outputs subject bounding boxes (bboxes) and 27 keypoints. Both the bounding boxes and the keypoints are generated with their respective confidence scores to ensure tracking reliability and data integrity. This native integration provides immediate zero-shot inference capabilities, effectively circumventing the labor-intensive requirement of manual frame annotation and iterative model training typically required by foundational tracking models, empowering researchers to rapidly extract complex behavioral motifs directly from raw video.

### Automated Data Format Conversion and High-Throughput Visualization

The conversion of tracking coordinates of animal into biologically meaningful, publication-ready figures often depends on a combination of disparate software tools. To address this bottleneck and maximize accessibility and user experience, we engineered a data format conversion module capable of universal parsing (Fig. 2 middle). This module autonomously detects legacy tracking data or outputs exported from other tracking systems (such as CSV or Excel files) and maps them into a canonical coordinate schema fully compatible with DeepLabCut. We explicitly adopted the DeepLabCut standard—the most widely utilized open-source tracking framework in ethology—to leverage its expansive community ecosystem and ensure seamless interoperability with existing downstream analytical tools.

Building upon this standardized data, we developed integrated functionalities for automated data visualization. For spatial analysis in 2D top-down assays, EthoClaw supports the programmatic generation of trajectory heatmaps and velocity heatmaps. Furthermore, for multi-cohort statistical comparisons, the platform utilizes the 2D xy-tracking coordinates to automatically compute standard kinematic metrics like total distance traveled based on predefined mathematical formulas. Relying on these derived metrics, the engine renders violin plots, multidimensional radar charts, and cluster maps to facilitate the identification of latent behavioral sub-types. Crucially, an automated typesetting algorithm successfully aggregates these independent plots into coherent, perfectly aligned composite figures suitable for manuscript preparation and writing.

### LLM-driven Methodological Transparency and Knowledge Integration

Upon completing an analytical pipeline, the platform synthesizes a comprehensive visual and textual summary report—integrating the generated figures with analytical overviews—which is automatically saved locally for convenient review. Should researchers have inquiries regarding the underlying analytical process, they can directly consult EthoClaw’s conversational interface to modify calculation methods or instantly retrieve transparent explanations—including exact mathematical parameters, velocity thresholds, and statistical assumptions—providing the precise verbiage required for publication (Fig. 2 bottom).

Driven by an embedded LLM, EthoClaw provides automated literature summarization—an essential tool for researchers trying to keep pace with the rapidly expanding volume of scientific publications (Fig. 2 bottom). The Knowledge Integration pipeline accomplishes this through two main approaches. First, it directly parses locally stored PDF files, extracting key information to create structured, archived summaries. Second, the system conducts automated web searches, either on a set schedule or on demand, to retrieve and summarize relevant recent papers. By integrating these functions, EthoClaw maintains a continuously updated, personalized local knowledge base and reading log. This drastically reduces the tedious manual labor of literature management, directly improving research efficiency.

### Application: Automated Phenotyping in an Open Field Test Paradigm

To validate the operational efficiency of EthoClaw, we evaluated a standard Open Field Test (OFT) paradigm in black mice. The subjects were assigned to two distinct experimental groups (Group A and Group B, n=5 per group) and tested in a standard 50 × 50 cm white opaque open field arena. Behavioral sessions were recorded from a top-down perspective at a resolution of 640 × 360 pixels for a duration of 5 minutes per animal at a sampling rate of 30 Hz, yielding a total of 9,000 frames per video for downstream computational analysis (Fig. 3a).

**Fig. 3.**
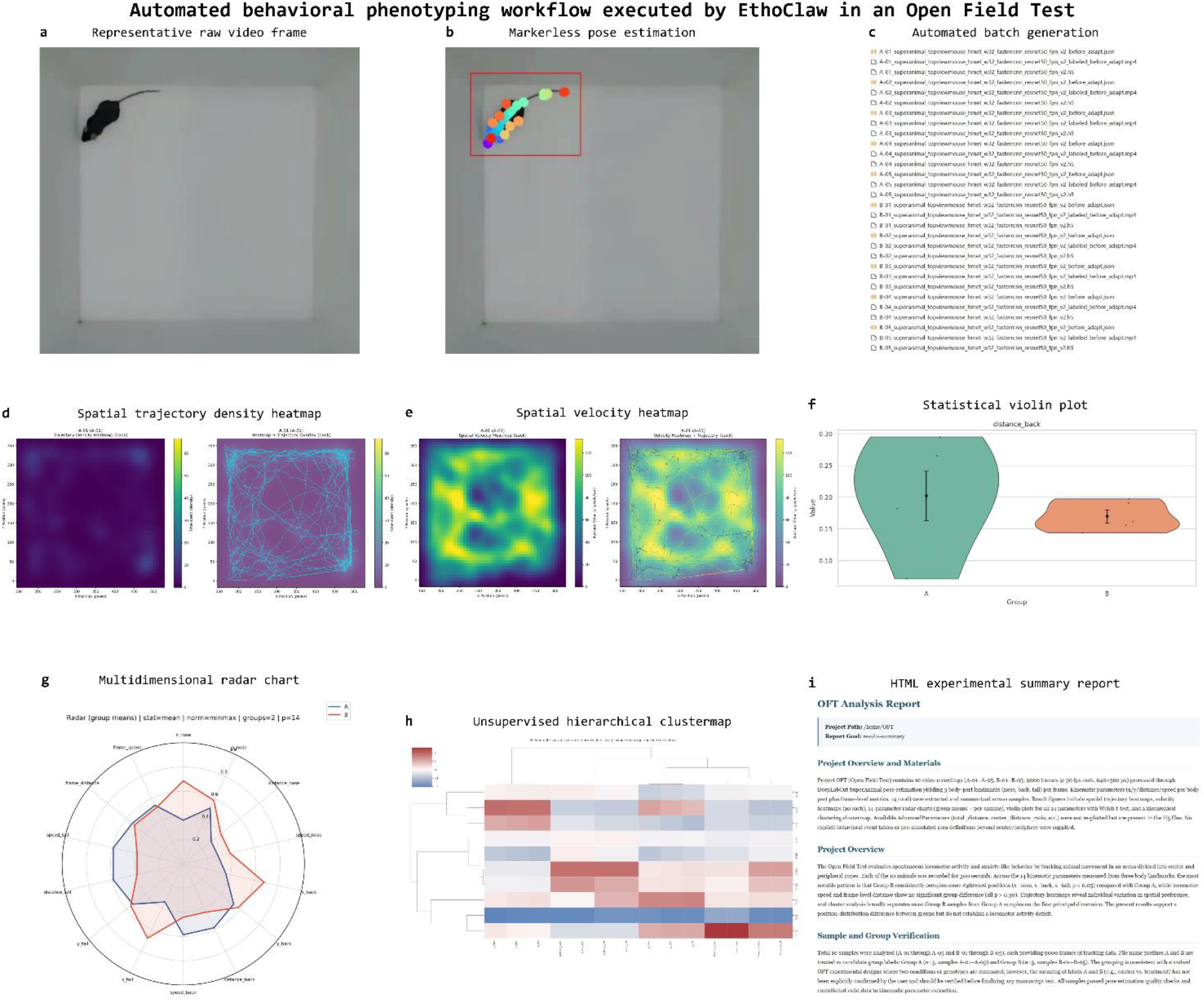
Automated behavioral phenotyping workflow executed by EthoClaw in an Open Field Test. **a**, Representative raw video frame capturing a subject traversing the open field arena. **b**, Markerless pose estimation utilizing the SuperAnimal deep learning model, successfully extracting 27 distinct anatomical keypoints per frame. **c**, Automated batch generation of standardized tracking data files formatted for seamless downstream processing. **d**, Spatial trajectory density heatmap illustrating the cumulative locomotor path and regional preference of a single representative subject. **e**, Spatial velocity heatmap demonstrating the instantaneous movement speed correlated with specific arena locations. **f**, Statistical violin plot visualizing the comparison of back-node travel distance between experimental Group A and Group B, featuring overlaid raw data points and central tendency markers. **g**, Multidimensional radar chart synthesizing the comprehensive kinematic profiles of both groups utilizing normalized mean values across 14 distinct behavioral parameters. **h**, Unsupervised hierarchical clustermap integrating all extracted kinematic features, demonstrating the data-driven categorization of individual samples and parameter covariation. **i**, Excerpt from the autonomously generated HTML/PDF experimental summary report, structurally aggregating project metadata, statistical inferences, and high-resolution visualizations.

The entire analytical workflow was orchestrated conversationally via the Lark messaging application, utilizing the built-in Channel function of the OpenClaw framework to communicate with EthoClaw, which was deployed locally on a high-performance workstation running Ubuntu 24.04 LTS and equipped with an NVIDIA RTX 4080 Super GPU. The analysis was initiated using the ethoclaw-animal-pose-estimation skill, which autonomously invoked the SuperAnimal model. By employing multi-process parallelization across all experimental videos, we rapidly extracted 27 two-dimensional anatomical keypoints per frame (Fig. 3b). However, rather than processing this dense coordinate dataset, we utilized a streamlined set of spatial coordinates previously obtained from an alternative pose estimation model to significantly accelerate downstream kinematic parameter extraction and high-throughput graphical rendering (Fig. 3c). This focused dataset explicitly comprised only three primary anatomical nodes: the nose, the back, and the tail. Utilizing these refined, computationally lightweight spatial coordinates, the ethoclaw-kinematic-parameter-generator skill was deployed to autonomously compute a comprehensive suite of kinematic parameters. Subsequently, the ethoclaw-trajectory-velocity-heatmap-generate skill was executed to render spatial trajectory and velocity heatmaps (Fig. 3d, e). To conduct rigorous multidimensional comparisons based on these derived parameters, we applied a sequence of specific skills—ethoclaw-multiparameter-violin-stats-generate, ethoclaw-multiparameter-clustermap-generate, and ethoclaw-multiparameter-radar-generate—seamlessly producing between-group statistical violin plots, multidimensional cluster maps, and radar charts (Fig. 3f-h). Following these quantitative analyses, the ethoclaw-analysis-report skill synthesized a detailed experimental summary report detailing the results (Fig. 3i). Finally, post-analysis knowledge integration was achieved by deploying the ethoclaw-daily-paper skill to search for the latest methodological advancements in behavioral analysis tools, automatically archiving the retrieved literature into local research logs.

The entire computational pipeline executed successfully without requiring any intermediate manual intervention. All invoked skills operated strictly as anticipated, reliably rendering the predefined high-resolution visualizations, multidimensional statistical plots, and the comprehensive HTML/PDF summary report. Furthermore, the automated literature retrieval functioned seamlessly, confirming EthoClaw’s robust end-to-end capability in facilitating highly integrated, efficient, and reproducible behavioral phenotyping.

## Discussion

EthoClaw addresses critical infrastructure bottlenecks in modern behavioral neuroscience by unifying fragmented analytical pipelines. By encapsulating raw video pose estimation, data wrangling, statistical validation, and report generation within a single locally deployed framework, EthoClaw eliminates the friction inherent in transitioning between disparate software environments. The integration of an API-driven LLM orchestration layer coupled with a local computational backend represents a highly pragmatic architectural paradigm; it democratizes access to state-of-the-art machine learning models via natural language while decisively circumventing the prohibitive bandwidth and latency bottlenecks associated with purely cloud-based video processing systems.

The introduction of a dual-mode tracking architecture further enhances the platform’s utility. By providing both ultra-fast, zero-annotation image processing and highly granular SuperAnimal pose estimation, EthoClaw affords researchers the flexibility to tailor the computational intensity to the specific demands of their experimental paradigms. Furthermore, aligning the Data Harmonization Module with DeepLabCut standards ensures that EthoClaw operates as an integrated bridge within the open-source community, preventing vendor lock-in and allowing users to leverage a vast ecosystem of downstream analytical tools.

Perhaps the most significant contribution of EthoClaw is its structural commitment to Open Science and experimental reproducibility. The historical reliance on undocumented custom scripts has severely hampered the reproducibility of behavioral phenotypes. EthoClaw’s automated generation of explicitly detailed “Methods” texts ensures that all mathematical thresholds and statistical assumptions are rigorously documented. This transparency empowers peer reviewers and future researchers to precisely replicate the computational conditions of published studies.

While EthoClaw exhibits exceptional performance in 2D top-down tracking scenarios, its deep learning optimizations are presently focused on single-animal paradigms, specifically calibrated for common models such as black mice. Subsequent development iterations will prioritize expanding the core computer vision engine to support complex 3D tracking environments and multi-animal assays, necessitating advanced identity maintenance algorithms. As an open-source project, we invite the computational biology community to contribute specialized visualization modalities and integrate diverse foundational tracking models via ClawHub or EthoClaw.

## Methods

### Dual-Mode Animal Object Detection and Pose Estimation

Raw experimental video streams are natively decoded via OpenCV, employing a generator pattern for sequential frame fetching to optimize memory utilization during extended, high-framerate behavioral sessions. To accommodate varying experimental demands—ranging from rapid, low-overhead spatial tracking to granular, high-dimensional postural analysis—EthoClaw deploys a flexible, dual-mode tracking architecture driven by specific analytical skills:

Object Detection via the ethoclaw-animal-grounding skill. Optimized for high-throughput, high-contrast paradigms such as tracking black mice on a light background, this highly CPU-efficient, zero-shot pipeline first applies grayscale conversion and 5×5 Gaussian smoothing to mitigate high-frequency noise and establish a standardized visual baseline. The smoothed frames then undergo inverse binary thresholding (pixel intensity ≤ 80) to isolate the animal’s silhouette. A subsequent morphological opening operation (5×5 kernel) eliminates micro-noise and background artifacts. The algorithm extracts external spatial contours, classifies the largest continuous region exceeding 300 pixels as the subject, and dynamically computes its geometric centroid (x_t, y_t) using spatial moments (m10/m00 and m01/m00). To preserve absolute temporal alignment during occlusions, missing contours are recorded as NaN coordinates with a 0.0 likelihood, while successful detections receive a pseudo-likelihood of 1.0. Finally, this mode autonomously generates a diagnostic video with centroid overlays for user validation and exports the temporal coordinates into a DeepLabCut-compatible multi-index schema (HDF5/CSV) for downstream harmonization.

Deep Learning Pose Estimation via the ethoclaw-animal-pose-estimation skill. For high-dimensional, markerless multi-node postural analysis, EthoClaw natively integrates DeepLabCut SuperAnimal pre-trained models. To overcome the computational latencies typically associated with deep neural network inference on high-resolution videos, EthoClaw implements a robust multi-process parallelization strategy. This architecture asynchronously distributes frame preprocessing tasks across available CPU cores before batching and loading tensors onto the GPU. For top-down mouse paradigms, the system deploys the superanimal_topviewmouse model, which is specifically trained to extract 27 distinct anatomical keypoints. These encompass the head and neck region (8 points: nose, left_ear, right_ear, left_ear_tip, right_ear_tip, left_eye, right_eye, head_midpoint), the torso midline and body (8 points: neck, mid_back, mouse_center, mid_backend, mid_backend2, mid_backend3, left_midside, right_midside), major limb joints (6 points: left_shoulder, right_shoulder, left_hip, right_hip, tail_base, tail_end), and detailed tail segments (5 points: tail1, tail2, tail3, tail4, tail5). Animal localization is initially performed using a Faster R-CNN detector with a ResNet-50 FPN v2 backbone, utilizing a strict bounding box detection threshold of 0.9. The localized batched tensors are then propagated through a High-Resolution Network (HRNet_w32) architecture for pose estimation. Unlike traditional convolutional networks that heavily downsample images, HRNet maintains high-resolution representations throughout the forward pass, ensuring the capture of fine-grained anatomical details necessary for extracting these distinct keypoints. Inference is executed via PyTorch, effectively leveraging hardware accelerators such as CUDA or MPS. Sub-pixel spatial coordinates (x_t, y_t) are derived from 2D probability heatmaps via soft-argmax operations. Concurrently, the network assigns a predictive confidence score (p between 0 and 1) to each identified keypoint. Operating with a pseudo-label threshold of 0.1, EthoClaw enforces rigorous downstream data integrity through a stringent confidence gating mechanism: any spatial coordinate predicted with a probability p < 0.8 is automatically nullified and encoded as a NaN value. The resulting high-fidelity coordinate datasets—structured with multi-index columns for scorer, individuals, bodyparts, and coords—are exported in HDF5 and CSV formats, and subsequently passed to the signal processing module for rigorous algorithmic interpolation and smoothing.

### Data Format Conversion

The Data Format Conversion Module, driven by the ethoclaw-normalize-tabular skill, processes raw coordinate timeseries from legacy systems and diverse third-party tracking software outputs such as comma-separated values, Excel spreadsheets, and HDF5 files. Utilizing highly vectorized Pandas and NumPy operations, the module inspects and normalizes local data structures into analysis-ready formats. Specifically, the parsing engine employs regex-based heuristics to automatically detect wide-format pose columns and structurally pivot them into a canonical long-form coordinate schema comprising frame, bodypart, x, y, and confidence variables. Additionally, the system seamlessly parses multi-dimensional HDF5 pose tensors, extracting axis labels to map them into the identical standardized long-form structure. Column names are rigorously standardized utilizing snake case formatting, missing value markers are algorithmically sanitized, and explicit data provenance variables are appended to ensure full methodological traceability. To optimize memory utilization and downstream analytical efficiency, the harmonized datasets are exported into the highly compressed Parquet format alongside automatically generated JSON schemas and diagnostic reports, ensuring seamless interoperability with subsequent kinematic profiling modules.

### Kinematic Parameter Extraction

Following data format conversion, EthoClaw derives advanced movement metrics utilizing the ethoclaw-kinematic-parameter-generator skill. This module autonomously processes the smoothed two-dimensional skeleton data residing within the standardized HDF5 files to compute a comprehensive suite of kinematic parameters.

To accurately calculate temporal dynamics, the system first determines the exact video framerate. It preferentially reads this value from internal HDF5 metadata. If the metadata is absent, the skill automatically scans the local directory structure to locate the corresponding raw video file, subsequently extracting the precise framerate natively utilizing FFmpeg or an OpenCV fallback routine. If no video file is located, a default framerate of 30 frames per second is strictly applied. Simultaneously, the system retrieves spatial calibration metrics—specifically pixel-to-millimeter ratios—to accurately map raw pixel displacements into absolute physical distances.

The kinematic extraction algorithm processes the coordinate arrays using highly vectorized NumPy operations. For every tracked anatomical keypoint, the module calculates the instantaneous physical coordinates converted to centimeters, the inter-frame Euclidean displacement, and the instantaneous spatial velocity. Furthermore, to evaluate whole-body locomotion, the system dynamically computes the geometric mean center across all detected body parts, deriving a global frame distance and global frame speed. The resulting multidimensional kinematic dataset is written directly back into the source HDF5 file under a dedicated parameter group. This in-place data integration ensures that raw coordinates, interpolation metadata, and derived kinematic features remain tightly coupled within a single portable and fully reproducible file architecture.

### Statistics and High-Dimensional Data Visualization

To facilitate comprehensive statistical evaluation, holistic behavioral profiling, and unbiased pattern discovery, EthoClaw employs a suite of specialized analytical skills that operate directly on the standardized HDF5 datasets. Engineered to seamlessly bridge the gap between high-dimensional kinematic data and publication-ready visual insights, these modules autonomously execute temporal data aggregation, systematic normalization, and rigorous hypothesis testing. Underpinned by core computational libraries including Pandas, SciPy, StatsModels, Scikit-learn, and Seaborn, the visualization and statistical pipeline is explicitly divided into three primary modules tailored to distinct analytical objectives:

Inferential Statistics and Distribution Visualization via the ethoclaw-multiparameter-violin-stats-generate skill. To rigorously evaluate group-level differences across extracted kinematic parameters, this module automates the inferential statistical pipeline directly from the standardized HDF5 files. During batch processing, temporal kinematic data are aggregated into per-sample scalar values, such as the temporal mean across all frames, to construct a unified feature matrix. The module processes statistical evaluations based on the experimental design and configurable parameters. For two-group comparisons, the system employs Welch’s t-test—ensuring robustness against unequal variances—or the non-parametric Mann-Whitney U test. For paradigms comprising three or more groups, the pipeline utilizes One-way Analysis of Variance or the non-parametric Kruskal-Wallis test. Crucially, if the overall multi-group test yields statistical significance, the system autonomously executes post-hoc pairwise comparisons and applies the Holm-Bonferroni method to rigorously correct p-values for multiple testing. The comprehensive statistical outcomes are seamlessly integrated into high-resolution violin plots rendered via the Seaborn library. These visualizations provide transparent representations of the underlying data distributions by explicitly overlaying raw data points with jitter, central tendency error bars denoting the mean and standard error of the mean, and automated significance brackets that display the adjusted p-values directly above the compared experimental cohorts.

Multivariate Behavioral Profiling via the ethoclaw-multiparameter-radar-generate skill. To construct a holistic visualization of the multivariate behavioral phenotype, EthoClaw utilizes this module to produce publication-ready radar charts. The system first extracts temporal kinematic data from the HDF5 files and aggregates the frame-by-frame timeseries into a single representative scalar per parameter for each sample, utilizing central tendency measures such as the mean, median, or maximum. To account for distinct physical units and scales across different kinematic features, the aggregated sample matrices are systematically normalized using either min-max scaling or Z-score standardization. Furthermore, the system autonomously infers experimental group assignments directly from the file nomenclature via regular expression parsing. For analytical flexibility, the module can generate individual multidimensional profiles for each specific sample, or compute and plot group-level mean polygons on a unified comparative radar chart. These visualizations are rendered employing predefined aesthetic presets inspired by top-tier scientific journals, generating high-resolution graphics that enable researchers to rapidly discern simultaneous multivariate behavioral shifts across different experimental cohorts.

Hierarchical Clustering and Clustermap Visualization via the ethoclaw-multiparameter-clustermap-generate skill. To objectively uncover latent behavioral sub-types and profile individual movement motifs without a priori labeling bias, this module executes an automated hierarchical clustering pipeline. First, the temporal sequence of every kinematic parameter within a sample is mathematically collapsed into a single representative scalar value—utilizing either the mean, median, or maximum operator—to construct a comprehensive feature matrix where rows represent independent samples and columns represent distinct kinematic parameters. Because parameters vary fundamentally in scale, the module systematically applies a column-wise Z-score normalization across all samples. The standardized feature matrix then undergoes agglomerative hierarchical clustering on both the sample and parameter axes. The algorithm computes linkages utilizing flexible distance metrics such as Euclidean or correlation distances, paired with linkage methods including average, complete, Ward, or single linkage. Finally, the module renders high-resolution, publication-ready clustermaps comprising a central heatmap flanked by respective dendrograms. To ensure optimal data presentation, the system includes predefined aesthetic presets inspired by top-tier journals, which automatically configure diverging colormaps, linewidths, and font scaling to deliver visually rigorous representations of parameter covariation and sample clustering.

### Experimental Summary Report Generation

EthoClaw employs a lightweight three stage workflow to automatically compile standalone Markdown and HTML/PDF reports from analytical results. The system first executes an automated project scan to catalog experimental artifacts including skeleton coordinate data, statistical tables and figure galleries. This module applies regular expression parsing to file nomenclature to infer experimental groupings and behavioral paradigms while extracting mathematical features from raw tracking files to compute trajectory summaries. These facts are centralized into a JSON manifest file outlining the project evidence. The large language model then reads this manifest and populates specific section bodies including sample validation, statistical insights and result interpretations. The agent synthesizes numerical trends and visual patterns while adhering to analytical guardrails to separate observed empirical facts from unverified mechanistic conclusions.

A dedicated rendering engine subsequently processes the populated manifest. Utilizing a custom Python based templating system the module assembles a comprehensive Markdown report and compiles it into a standalone HTML/PDF document. To ensure document portability, the rendering pipeline leverages image processing libraries to optimize and encode all visual assets into base64 data URIs. Embedding all experimental figures directly within a single HTML/PDF file eliminates the risk of broken image links and delivers an output suitable for laboratory archiving and scientific sharing.

### Knowledge Integration and PDF Document Summarization

EthoClaw incorporates a document processing module to ingest and parse local academic publications and technical reports. The engine employs a dual path extraction workflow to handle diverse document formats. The system utilizes command line utilities including pdfinfo and pdftotext to extract metadata and layout preserved text. For documents with complex formatting or scanned content the module invokes pdftoppm to render high resolution page images. This extraction phase produces an analysis bundle containing a centralized JSON manifest paired with a raw text transcript and sequential page previews.

Following extraction the agent evaluates the generated manifest and textual data. If the extracted text appears fragmented or sparse the system transitions to visual inspection of the rendered page images to interpret table structures and scientific figures. The platform then synthesizes the document content into structured deliverables based on user confirmation. The system generates standardized research logs and summaries that categorize core arguments along with methodological evidence and experimental findings. The agent writes these outputs directly into the local workspace as Markdown/PDF files to establish a persistent and searchable research library.

### Online Literature Search and Summarization

To facilitate continuous tracking of methodological advancements EthoClaw implements an automated literature retrieval module targeting external databases including PubMed and arXiv. The system parses a local configuration file to construct database specific query strings and establish inclusion and exclusion keyword lists relevant to neuroethology and behavioral neuroscience. The execution pipeline queries the respective application programming interfaces to retrieve recent publications within a specified temporal window. The module extracts metadata including titles abstracts publication dates and author lists while applying a weighted scoring algorithm that prioritizes peer reviewed PubMed articles over arXiv preprints and rewards keyword density alongside relevant subject categories.

Following data extraction the system aggregates the retrieved records and performs cross database deduplication using unique identifiers such as digital object identifiers or cryptographic hashes of the article titles. This process generates a centralized metadata payload and a consolidated candidate pool sorted by mathematical relevance. To optimize computational resources during evaluation the platform employs a single session workflow that initially exposes only the publication titles to the large language model. The agent evaluates this title surface against the configured research priorities to autonomously select the top five candidate papers circumventing the need to process the entire abstract corpus during the preliminary ranking phase.

Upon finalizing the target selection the agent accesses the complete abstracts of the five chosen publications to synthesize a comprehensive research digest. The framework dictates a strict output structure where the model writes a continuous explanation encompassing background context experimental methods main findings scientific significance and identified limitations. The system instructs the agent to preserve concrete experimental variables including target species brain regions behavioral paradigms and neural recording techniques while explicitly stating uncertainties when the abstract lacks sufficient evidentiary support. The final deliverable is rendered directly into a standalone Markdown document within the local workspace providing researchers with a structured daily literature overview without invoking external text summarization application programming interfaces.

### Performance Optimization

EthoClaw incorporates multiple optimization strategies at the core execution layer to maintain computational efficiency during continuous analytical workflows. The platform implements an intent driven lazy loading mechanism for tool schemas. The embedded engine evaluates user queries against predefined keyword heuristics to categorize tasks and dynamically load only the necessary tool profiles. This conditional loading logic applies to primary sessions as well as background processes and scheduled tasks directly reducing total token consumption and system memory overhead.

The framework addresses context window exhaustion through an automatic truncation protocol for oversized file extraction and programmatic outputs. When a tool result exceeds a specific character limit the platform truncates the central portion of the payload while retaining the initial and final text segments. This intervention prevents memory overflow failures and preserves the structural data required by the language model for accurate analytical interpretation.

Operational latency is further reduced via a specialized prompt configuration that eliminates redundant system instructions and streamlines the overall context payload. The execution engine propagates this optimized prompt mode to all subordinate agents to ensure computational consistency. The platform also improves longitudinal memory management by expanding the retention capacity for exact conversational history and executing an automated state refresh following context compaction. These combined modifications ensure the autonomous agent maintains its primary research objectives over prolonged sessions without degrading inference speed.

## Data and Code Availability

EthoClaw is an open-source project licensed under the MIT License. The source code, documentation, and demo results are all freely available in the project repository: [https://github.com/penciler-star/EthoClaw].

